# A Novel human IL-23A Overexpressing Mouse Model of Systemic Lupus Erythematosus

**DOI:** 10.1101/2023.11.08.566163

**Authors:** Eleni Christodoulou-Vafeiadou, Christina Geka, Lida Iliopoulou, Lydia Ntari, Maria C. Denis, Niki Karagianni, George Kollias

## Abstract

**Objective:** Interleukin-23 (IL-23) is a crucial cytokine implicated in chronic inflammation and autoimmunity, associated with various diseases like psoriasis, psoriatic arthritis, and systemic lupus erythematosus (SLE). This study aimed to create and characterize a transgenic mouse model (TghIL23A) overexpressing human IL23A, providing a valuable tool for investigating the pathogenic role of hIL23A and evaluating the efficacy of anti-human-IL23A therapeutics.

**Methods:** TghIL23A mice were generated via microinjection of CBAxC57BL/6 zygotes with a fragment of the human IL23A gene, flanked by its 5’-regulatory sequences and the 3’UTR of human beta-globin. The TghIL23A pathology was assessed through hematological and biochemical analyses, cytokine and anti-nuclear antibody detection, histopathological examination of skin and renal tissues. The response to the anti-hIL23A therapeutic agent guselkumab, was evaluated in groups of 8 mixed-sex mice receiving subcutaneous treatment twice weekly for 10 weeks, using clinical, biomarker and histopathological readouts.

**Results:** TghIL23A mice exhibited interactions between hIL23A and mouse IL23/IL12p40, and developed a chronic multiorgan autoimmune disease marked by proteinuria, anti-dsDNA antibodies, severe inflammatory lesions in the skin and milder phenotypes in the kidneys and lungs. The TghIL23A pathological features exhibited significant similarities to those observed in human SLE patients.

**Conclusions:** We have generated and characterized a novel genetic mouse model of SLE, providing proof-of-concept for the etiopathogenic role of hIL-23A. This new model has a normal lifespan and integrates several characteristics of the human disease’s complexity and chronicity making it an attractive preclinical tool for studying IL23-dependent pathogenic mechanisms and assessing the efficacy of anti-hIL23A or modeled disease-related therapeutics.

## Introduction

Interleukin-23 (IL23) is a member of the IL12 cytokine family, composed of the IL23A (IL23p19) and the IL12/23B (IL12/23p40) subunits (1). It is primarily secreted by activated macrophages, dendritic cells (DCs), keratinocytes and other antigen-presenting cells, as well as by T lymphocytes and NK cells in peripheral tissues, such as the skin, intestinal mucosa, joints and lungs (2–4). While the IL23/IL17 axis has a well-documented protective role against bacterial and fungal infections (5), its dysregulation can lead to chronic inflammation and autoimmunity, contributing to the development of several diseases like psoriasis, psoriatic arthritis, inflammatory bowel disease, rheumatoid arthritis, multiple sclerosis and others (5). Due to its significant involvement in various pathogenic conditions, IL23 has emerged as a promising therapeutic target for chronic inflammatory and autoimmune diseases. Guselkumab, a monoclonal antibody for the treatment of moderate to severe plaque psoriasis, targeting IL23A and Ustekinumab, a monoclonal antibody for the treatment of Crohn’s disease, ulcerative colitis, plaque psoriasis and psoriatic arthritis, targeting the common p40 subunit of IL12 and IL23, are two of the currently approved biologics that target IL23, while additional biologics targeting IL23 are also under development.

One autoimmune disease where IL23 has recently been shown, both in human patients and mouse models, to play a crucial pathogenic role, is Systemic Lupus Erythematosus (SLE). SLE patients exhibit elevated IL23 levels in their sera and plasma, along with increased IL23R expression on T-cells (6–8). Conversely, IL23R deficiency has been found to protect the B6/*lpr* lupus-prone mouse model from lupus development (9) and treatment of the MRL/lpr genetic mouse model of lupus with anti-IL23A antibodies has shown to ameliorate lupus symptoms (10).

SLE is an autoimmune disease characterized by the loss of tolerance against nuclear autoantigens, resulting in complement and cytokine activation and the production of high levels of anti-single stranded and anti-double stranded autoantibodies (11–13). The disease is characterized by the presence of circulating immune complexes that deposit in tissues, leading to multiorgan damage, with kidneys and skin being primarily affected (13–16). Mouse models that replicate key features of SLE pathology have greatly contributed to the study of SLE etiopathogenesis and the identification and validation of potential therapeutic targets.

Among the commonly used mouse models for studying lupus pathology and for the preclinical evaluation of therapeutics, the NZB/W F1 model exhibits severe lupus-like phenotypes including splenomegaly, elevated levels of circulating anti-nuclear antibodies, immune complex-mediated nephritis leading to renal failure and premature death (17–19). The MRL/lpr mouse model, homozygous for the lymphoproliferation spontaneous mutation (Fas^lpr^), shows defective immune cell apoptosis, leading to the accumulation of double negative autoreactive (CD4^−^CD8^−^) B220+ T cells, the production of anti-dsDNA autoantibodies and the development of renal and skin pathologies (17–19). The BXSB/Yaa mouse model, a recombinant inbred strain, develops lymphoid hyperplasia, nephritis, high serum-titers of antinuclear antibodies and high-serum retroviral glycoprotein gp70 titers (18). Additionally, several mouse models based on single gene knockouts or transgenic expression of specific genes exhibit lupus-like phenotypes, providing insights into the involvement of these genes and functional pathways in SLE mechanisms (20). However, no mouse model fully recapitulating the complexity of human lupus pathology has been developed so far.

Considering the accumulating evidence on the pathogenic role of IL23, we generated TghIL23A transgenic mice expressing dysregulated human IL23A. We demonstrate that TghIL23A mice develop phenotypes that replicate major clinical, immunological and molecular features of human lupus erythematosus, making them a valid novel mouse model of lupus and a valuable preclinical tool for studying IL-23-dependent pathogenic mechanisms and evaluating therapeutics.

## Material and methods

### Generation of TghIL23A transgenic mice

A 2825 bp genomic DNA fragment containing 986 bp of the human *IL23A* 5’ regulatory sequences and the complete intron-exon sequences up to the stop codon was PCR amplified from the Bacterial Artificial Chromosome_CTD-3110H14 and it was further cloned upstream of a 779 bp long DNA fragment containing the 3’-untranslated and polyadenylation sequences of the human β-globin gene (21). This transgene was microinjected in (CBA x C57BL/6) F2 mouse zygotes and the transgene carriers were identified by PCR using the following primers F: GTCTTTGCCCATGGAGCAGC and R: GCCCTTCATAATATCCCCCA.

Pronuclear injection of the transgene into fertilized (C57BL/6JXCBA/J) F2 oocytes was performed in the BSRC Alexander Fleming Transgenesis Facility.

### Mice and in vivo studies

WT, TghIL23A, TghIL23A/IL12B-/-(22) and TghIL23A/Rag1-/-(23) mice were bred and maintained in a mixed CBA×C57BL/6J genetic background in the animal facilities of Biomedcode Hellas S.A. and BSRC Al. Fleming under specific pathogen-free conditions. Animals were housed in standard plastic cages with wood chip bedding, under an inverted 12:12-h light/dark cycle at a constant temperature of 22±2 °C and relative humidity of approximately 60%. Standard diet and water were provided *ad libitum*.

For *in vivo* studies, 10-week-old mice were distributed to sex-balanced groups of 8-10 animals and were treated subcutaneously twice weekly for 10 weeks with either saline or different doses of Guselkumab (Tremfya, Janssen-Cilag International NV). Mice were monitored regularly to record their weight and skin clinical scores while ear thickness was measured at the completion of the *in vivo* part of the study.

All animal experimentation was approved by the BSRC Al. Fleming Institutional Committee of Protocol Evaluation in conjunction with the Veterinary Service Management of the Hellenic Republic Prefecture of Attica according to all current European and national legislation and were performed in accordance with relevant guidelines and regulations (approved protocol No 466365/25.05.22).

### Skin clinical score

The TghIL23A skin phenotype was assessed based on a scoring scale adapted from Yang JQ et al. (24) with clinical scores representing the sum of 3 individual scores assessing 1) the extend of periocular fur loss: 0 (normal), 0.5 (mild), 1 (severe); 2) the extend of fur thinning and skin redness of the abdominal area: 0 (normal), 0.5 (mild), 1 (severe) and 3) the severity of the ventral neck and thorax area pathology: 0 (normal), 0.5 (mild lesions in the neck area), 1 (mild to moderate lesions extending towards the thorax) and 2 (severe extended skin lesions with serous exudates and crusts). Ear thickness was measured using an electronic caliper.

### Cytokine and autoantibody detection

Cytokines were measured in skin tissue lysates using the LEGENDplex™, Mouse Inflammation Panel (13-plex) according to manufacturer’s instructions. Serum levels of anti-dsDNA, anti-ssDNA antibodies and total IgG were assessed by ELISA (Alpha Diagnostic International; 5120, Alpha Diagnostic International; 5320 and Abcam; ab157719 respectively) according to the manufacturer’s instructions.

### Hematological and Urine analysis

Red blood cells (RBS), hematocrit (HCT) and hemoglobin levels were measured using the Mindray BC-5000Vet Hematology Analyzer. Urine protein concentrations were assessed either with the Beckman Coulter AU480 Clinical Chemistry Analyzer or using Multistix 10 SG (SIEMENS; 2300).

### Flow cytometry of skin samples

Skin samples collected from the ventral neck area of 4-5 mice were minced in Hank’s balanced solution (HBSS) and incubated at 4°C overnight with 1.5mg/ml dispase II (Roche; 25766800). The next day, samples were incubated at 37°C with 1mg/ml collagenase IV (SIGMA; C5138) and 10%FBS for 1.5h while shaking. Following this incubation, the samples were vortexed, filtered through a 70-μm cell strainer, washed with HBSS+10%FBS and centrifuged at 1200 rpm. The cell pellet was resuspended in FACS buffer and 1.5×10^6^ cells/sample were stained with the viability dye Zombie-NIR (Biolegend) and with specific antibodies for the detection of: CD4, CD8, B220, CD11b, Ly6G, F4/80, SinglecF (BioLegend) and analyzed by flow cytometry with BD FACS Canto II and BD FACSDIVA software.

### Histopathology and immunohistochemistry

Tissue paraffin sections were stained with H&E for histopathological evaluation or with biotinylated anti-mouse-IgG (Vector; BA-9200) for the detection of immune complex deposits and anti-human IL23A (OriGene; AM20386PU-N) for the detection of human IL23A protein in various tissues.

Skin and ear histopathological evaluation was performed in a blinded manner using a scoring scale adapted from Yang JQ et al (24). Scores are the sum of the individual scores of epidermis and dermis pathology. Epidermis pathology is characterized by hyper/orthokeratosis and acanthosis leading to tissue destruction and necrosis assessed in a scale from 0-4 and dermis pathology includes immune cell infiltration in the different dermis layers assessed in a scale ranging from 0-3.

Lung histopathological evaluation was also performed in a blinded manner using a scoring scale adapted from Bell et al. (25). Scores are the sum of the individual scores of peribronchial and perivascular inflammation (scale 0-3), along with interstitial inflammation (scale 0-3).

### RNA isolation and PCR analysis

Tissue RNA was isolated with Trizol (ThermoFisher scientific; 15596026) and reverse transcribed with M-MLV Reverse Transcriptase (Promega; M1705). The expression of human IL23A mRNA and β2m was assessed by PCR using specific primers (hIL23AF: GTCTTTGCCCATGGAGCAGC; hIL23AR: GCCCTTCATAATATCCCCCA; mβ2mF: TTCTGGTGCTTGTCTCACTGA; mβ2mR: CAGTATGTTCGGCTTCCCATTC).

### Statistical analysis

Data are presented as mean ± SEM, and Student’s t*-test* was used for the evaluation of statistical significance, with *P* values < 0.05 being considered statistically significant. More specifically, Normality and Lognormality Tests were performed, choosing Kolmogorov Smyrnov, Shapiro-Wilk and D’Agostino and Pearson normal Gaussian distribution. If sample values passed all three tests, unpaired parametric Student’s t*-test* was chosen. If not, nonparametric Mann Whitney test was used. Analysis was performed using GraphPad Prism V.10.

## Results

### 3.1 Generation of human IL23A transgenic mice

To examine the pathogenic role of IL23A, we generated TghIL23A transgenic mice. The transgene used contained a fragment of the human IL23A gene locus comprised of 5’ regulatory sequences sufficient to confer regulated transcription (26) of the complete intron-exon sequences that followed. The fragment was cloned upstream of a non-regulated 3’ UTR to enable the abolishment of the posttranscriptional regulation of IL23A and its altered expression, aiming to increase the risk of development of a pathogenic condition (27) (**Fig. 1A**). The TghIL23A transgenic mice generated were maintained in CBAxC57BL/6 background and the expression of the human IL23A transgene was confirmed by PCR in the skin, lymph nodes, spleen, lungs and kidneys (**Fig. 1**Β). The human IL23A protein, was also detected by immunohistochemistry in the skin, showing a diffuse positive staining in the dermis, in the cortex of the lymph nodes, in the red pulp of the spleen, in the peribronchial and in perivascular areas as well as in the interstitium of the lungs and in the glomeruli of the kidneys (**Fig. 1C**).

**Figure 1:**
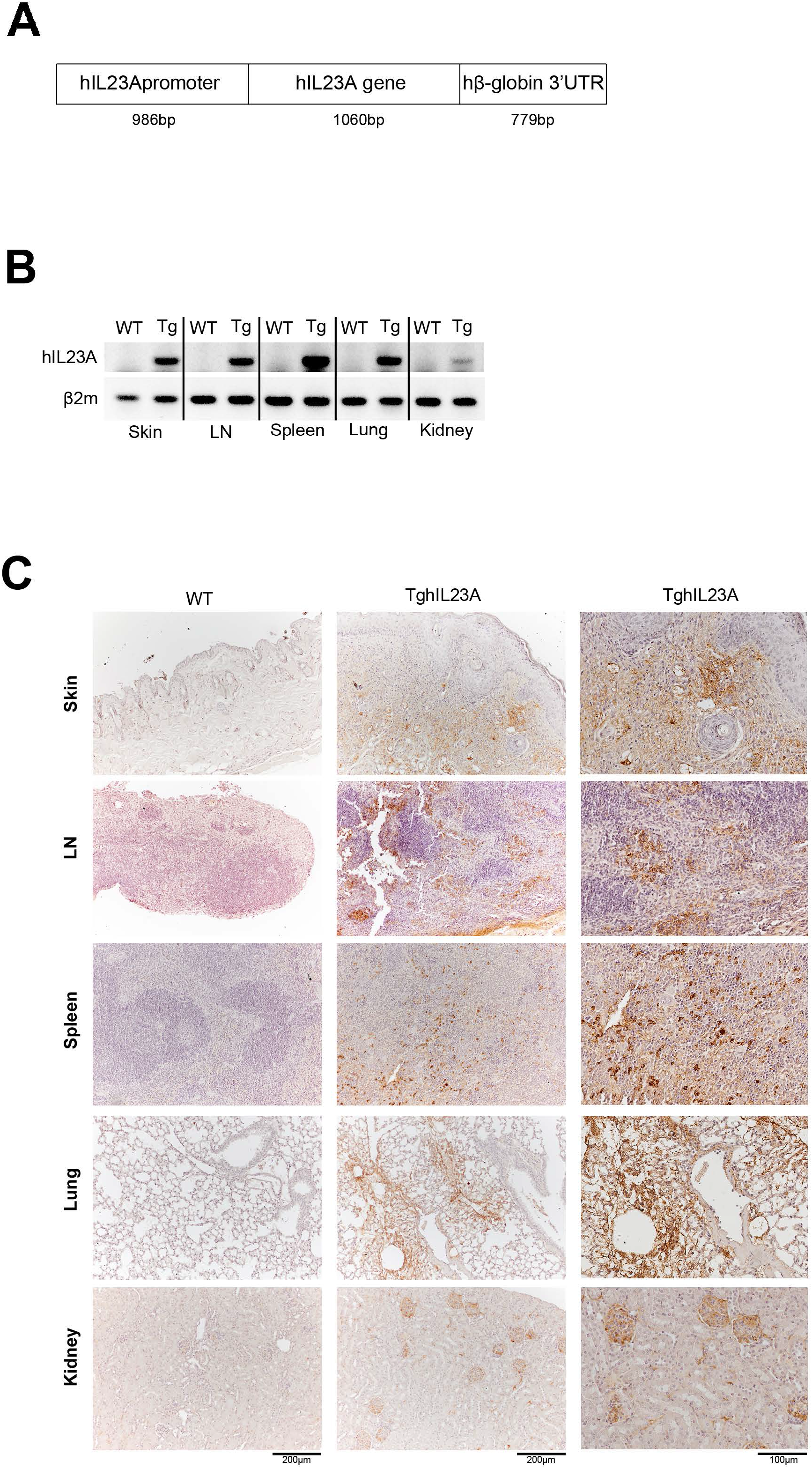
Generation of TghIL23A mice and hIL23A expression. **(A)** Schematic representation of the human IL23A transgene**. (B)** human IL23A mRNA detected by PCR and **(C)** human IL23A protein detected by immunohistochemistry in the skin, cervical lymph nodes, spleen, lung and kidney of TghIL23A mice. Scale bar 200μm and 100μm.

### 3.2 Human IL23A expression leads to a spontaneous and progressive skin pathology with immune system involvement

TghIL23A mice are fertile with normal life-span and exhibit normal body weight gain until the age of 7 months where they start to show a progressive body weight loss (**Supplementary Fig. 1**). At 4 weeks of age, the TghIL23A mice start developing clinical symptoms of skin pathology, first manifesting as fur thinning in the periocular and abdominal areas (**Fig. 2A, i**&iv). As the disease progresses, the clinical symptoms become more severe and include the thickening of the ear pinna (**Fig. 2A**, v-vi) as well as the extensive loss of fur and the appearance of skin lesions with serous exudate production and crust formation in the rostrum and the ventral neck/ upper thorax (**Fig. 2A**, ii-iii).

**Figure 2:**
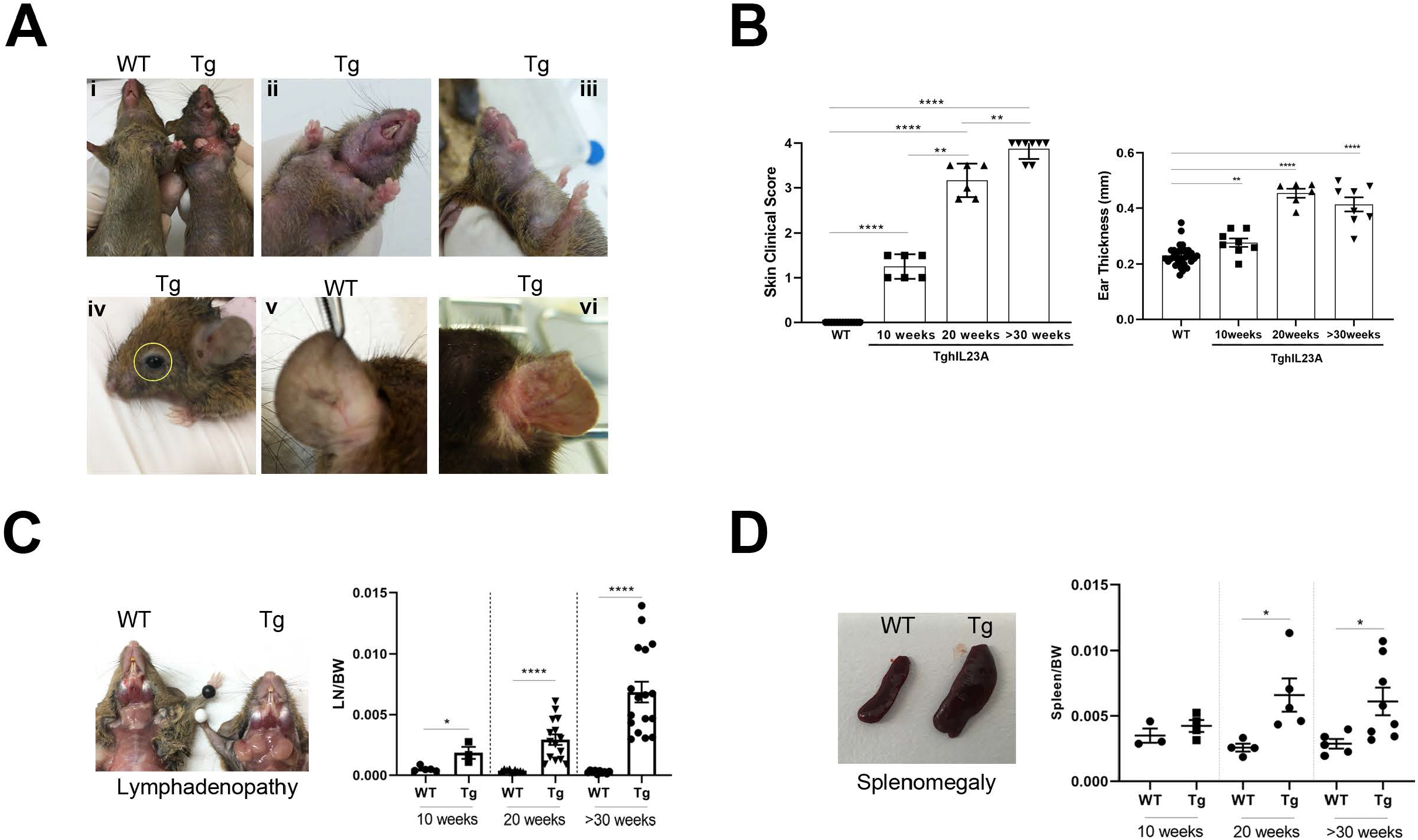
Clinical features of dermatitis, splenomegaly and lymphadenopathy in the TghIL23A mice. **(A)** Macroscopic images of 30-week-old TghIL23A (Tg) and wild type (wt) littermate mice depicting clinical features of fur thinning and skin lesions in the ventral-thoracic (i-iii) and periocular areas (iv) as well as in the ears of the transgenic mice (v-vi). **(B)** Clinical assessment and ear thickness measurements show an age-dependent increased severity of the TghIL23A skin pathology. The assessment of the lymph node and spleen to body weight ratios show the development of a TghIL23A age-dependent lymphadenopathy **(C)** and splenomegaly **(D)**. Data are presented as mean ± SEM; \**P* < 0.05; \*\**P* < 0.005; \*\*\**P* < 0.0002; \*\*\*\**P* < 0.0001.

The progression of the clinical phenotype was assessed using a scoring scale of four points taking into account the features of the pathology developing in the periocular, abdominal, ventral neck and thorax areas. By the age of 10 weeks all the animals developed mild skin pathology, that reached maximum score by 30 weeks of age (**Fig. 2B**). The clinical signs also include the involvement of the immune compartment as indicated by the hyperplastic phenotype observed in the draining lymph nodes of the affected ventral neck area (F**ig. 2C**), as well as the greatly enlarged spleens of 5-7-month-old mice (**Fig. 2D**).

### 3.3 Human IL23A expressing mice develop autoimmune lupus pathology

To further assess the clinical profile of the TghIL23A mice, we performed hematological and biochemical analysis of the blood and urine of 10-20-week-old mice. Analysis of the blood of 20-week-old TghIL23A mice showed significantly reduced red blood cell counts (RBC) as well as reduced hematocrit (HCT) and hemoglobin (HGB) levels, indicating an anemic phenotype (**Fig. 3A**). Moreover, urine analysis showed that, as early as 10 weeks of age, the TghIL23A mice exhibit proteinuria indicated by the increased levels of albumin and creatinine, suggesting renal dysfunction (**Fig. 3B**).

**Figure 3:**
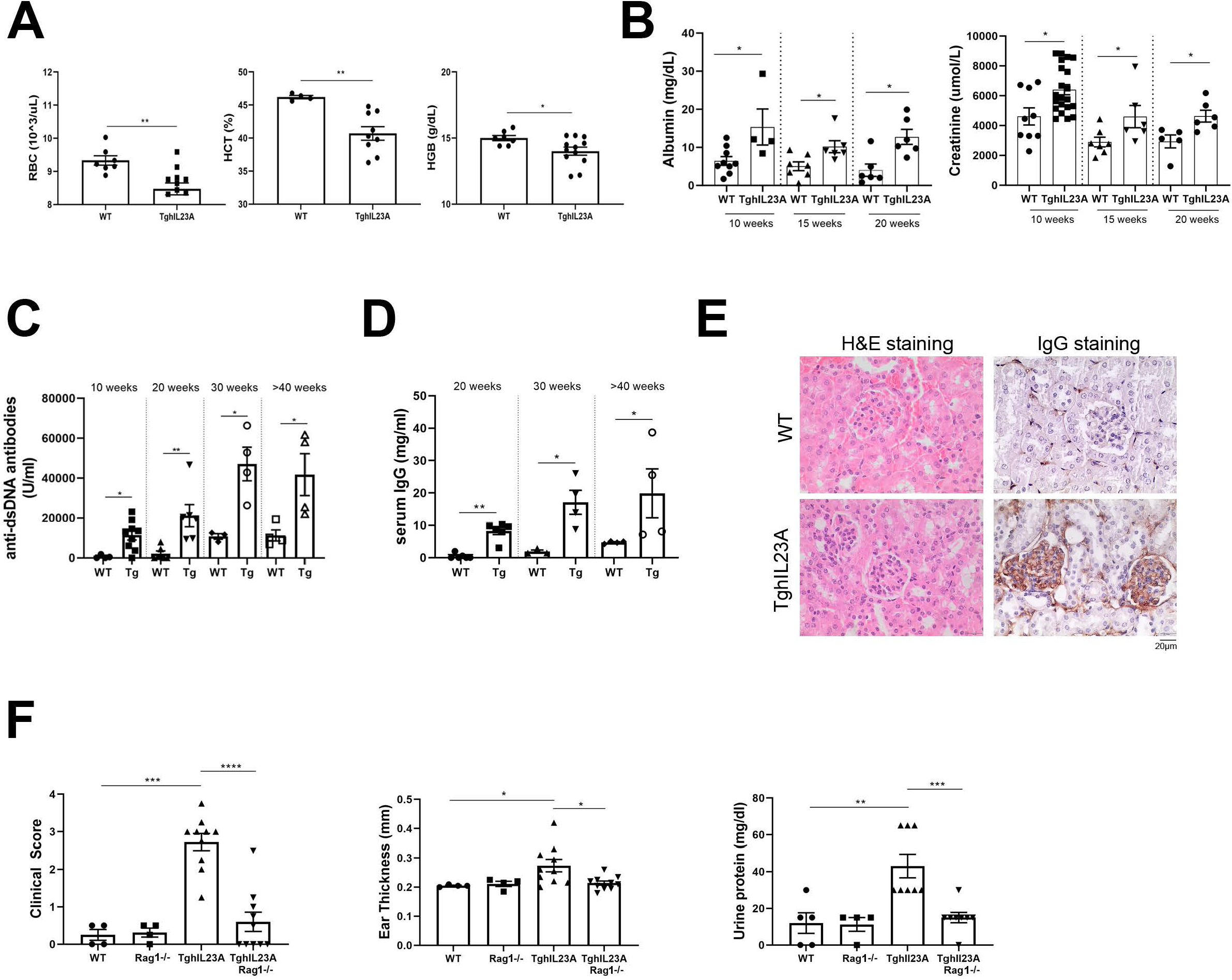
TghIL23A mice develop an autoimmune lupus-like pathology. 20-week-old TghIL23A mice show **(A)** signs of hemolytic anemia evident by the Reduced Red Blood Cell (RBC) counts, Hematocrit (HCT) and Hemoglobin (HGB) levels, **(B)** mild proteinuria evident by the increased Albumin and Creatinine urine levels, **(C)** increased circulating levels of anti-double-stranded DNA (anti-dsDNA) antibodies and **(D)** total IgG and **(E)** mild mesangial hypercellularity and IgG deposits in their glomeruli. Scale bar 20μm. **(F)** The TghIL23A clinical pathology features of 20-week-old mice are abolished following their crossing to Rag1-/-mice. Data are presented as mean ± SEM; \**P* < 0.05; \*\**P* < 0.005; \*\*\**P* < 0.0002; \*\*\*\**P* < 0.0001.

The finding of anemia, which is a common sign of chronic inflammatory conditions such as autoimmune pathologies and that of proteinuria, that implicates impaired renal function, suggested that the TghIL23A phenotype could possibly be linked with autoimmunity. To explore this possibility, we assessed key features of autoimmunity, including the presence of circulating autoantibodies as well as the T- and B-cell involvement. We therefore screened sera of wild type and TghIL23A mice at different ages for the detection of antinuclear antibodies (ANAs) and total immunoglobulin levels. From 10 weeks of age the sera of TghIL23A mice tested positive both for anti-single stranded (**Supplementary Fig. 2**) as well as anti-double stranded DNA antibodies (**Fig. 3C**). IgGs were significantly increased in circulation (**Fig. 3D**) and were deposited in the renal glomeruli, where also increased numbers of mesangial cells were detected (**Fig. 3E**). Crossing the TghIL23A mice to Rag1-/-mice resulted in complete abolishment of the skin pathology as well as of proteinuria (**Fig. 3F**), confirming the critical involvement of B and T lymphocytes.

Crossing the TghIL23A mice with the IL12B deficient mice (TghIL23A/Il12B-/-) resulted in the significant decrease of circulating anti ds-DNA antibodies (**Supplementary Fig. 3A**) and decreased proteinuria (**Supplementary Fig. 3B**) at 20-week-old mice. These data suggest that the pathology is driven by an IL23 dimer, composed of a human IL23A and a mouse IL12B subunit.

The simultaneous manifestation of inflammation in the skin, the impaired renal function and anemia together with the presence of antinuclear antibodies and the dependency of the pathology on T and B cells as well as on the expression of IL12B subunit, support that TghIL23A mice reproduce key features of systemic lupus autoimmunity (13,28–30) that are IL23-dependent.

### 3.4 TghIL23A mice develop cutaneous lupus with pulmonary manifestations

We next studied in greater detail the skin pathology and its connection with the autoimmune lupus developing in these mice. Histopathological analysis of H&E-stained sections of the affected skin of 20-30-week-old TghIL23A mice revealed the presence of vacuolar interface dermatitis (**Fig. 4A, i**), acanthosis of the epidermis with elongated rete ridges and orthokeratosis (**Fig. 4A, ii**), epidermolysis (**Fig. 4A, iii)** as well as accumulation of inflammatory cells (PMNs), ulcerations and necrosis in the dermis (**Fig. 4A, iv**). Similar, but slightly milder in severity, pathological features were observed in the ears of these mice that showed hyperkeratosis, acanthosis and accumulation of inflammatory cells (PMNs) in the subepidermal layer (**Supplementary Fig. 4A**).

**Figure 4:**
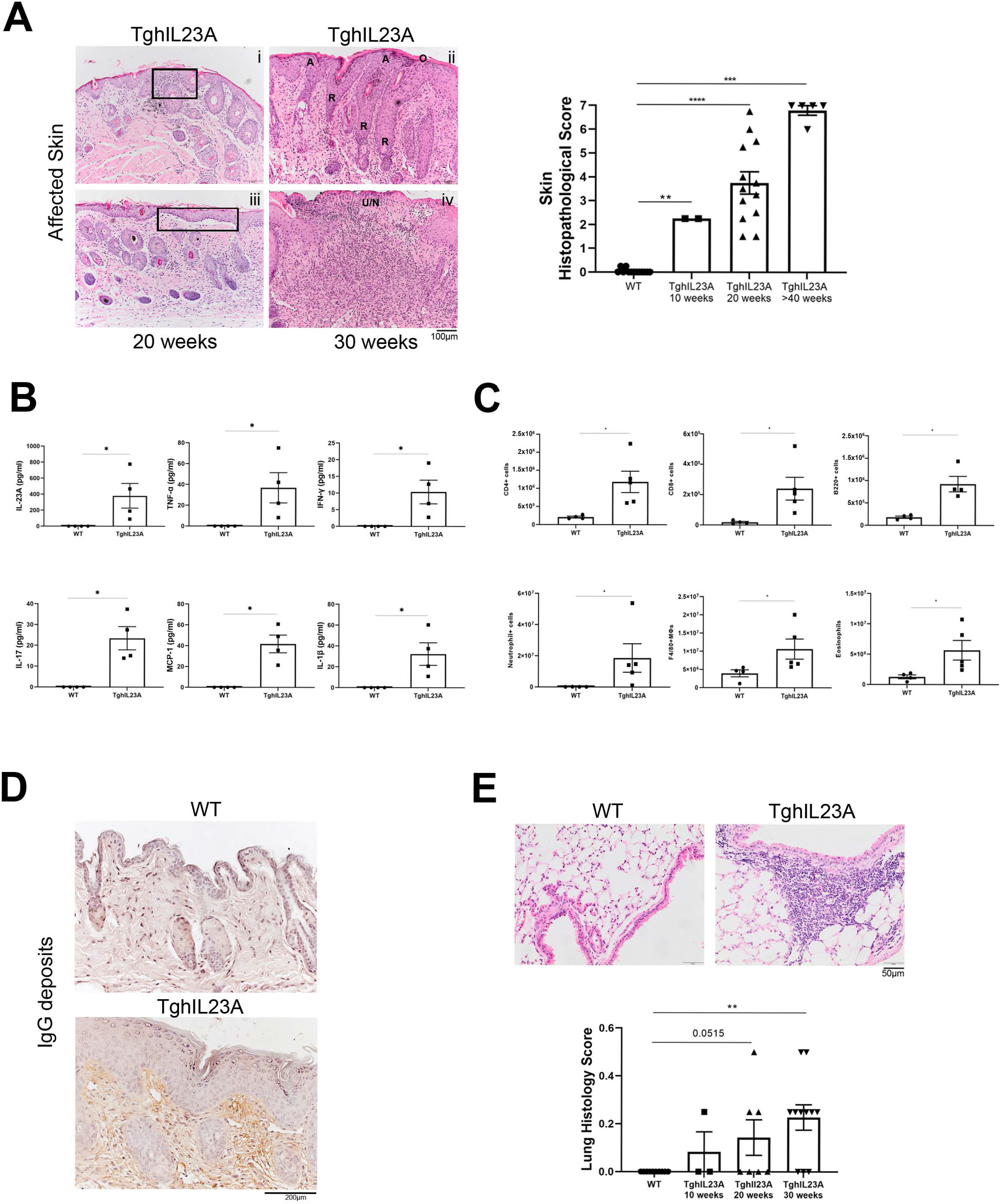
Histopathological analysis of the cutaneous and lung manifestations of the TghIL23A pathology. **(A)** H&E-stained paraffin sections showing vacuolar interface dermatitis (**i**), orthokeratosis (O), acanthosis (A), rete ridges formation (R) (**ii**), epidermolysis **(iii)** as well as accumulation of inflammatory cells (PMNs), ulcerations and necrosis (U/N) (**iv**) in the skin of TghIL23A mice. Scale bar 100μm. These pathological features become progressively more severe in older mice (graph). **(B)** The affected skin of 20-week-old TghIL23A mice also exhibit increased levels of inflammatory cytokines **(C)**, lymphoid and myeloid infiltrates as well as **(D)** IgG deposits. Scale bar 200um. **(E)** representative H&E-stained lung paraffin sections showing evidence of interstitial and peribronchial lung inflammation in 20-week-old TghIL23A mice that are also assessed for their severity histopathologically (graph). Scale bar 50μm. Data are presented as mean ± SEM; \**P* < 0.03; \*\**P* < 0.0095; \*\*\**P* =0.0001; \*\*\*\**P* < 0.0001.

The severity and progression of the skin pathology was monitored using a scoring scale of seven points assessing histopathological features present in the epidermis and dermis layers of the affected skin. In all the TghIL23A animals examined, mild skin and ear histological features were detected by the age of 10 weeks while scores reached maximum severity by the age of 40 weeks (**Fig. 4A** **and Supplementary Figure 4B**). Interestingly, similar to what we observed with the clinical features, the histopathological features in the skin were also abolished when the TghIL23A mice were crossed with Rag1-/-as well as with IL12B-/-mice (**Supplementary Fig. 5A, B**) highlighting again the dependency on the IL23 pathogenic pathway and the adaptive immune response.

Cytokine and chemokine multiplex analysis revealed that the local skin inflammatory milieu, contained increased levels of multiple inflammatory molecules such as IL23A, TNF-α, IFN-γ, IL-17A, MCP-1 and IL-1b (**Fig. 4B**). FACS analysis indicated the accumulation of inflammatory cells in the affected skin that included primarily CD4^+^ and CD8^+^ T cells, B220^+^ B cells, F4/80^+^ macrophages, neutrophils and eosinophils (**Fig. 4C**). In accordance with the accumulation of B cells in the skin of TghIL23A mice, extensive IgG deposits were detected in the upper dermis area (**Fig. 4D**).

Since lupus can affect multiple organs, we performed histopathological analysis of H&E-stained sections of the ileum, colon, heart and lungs, organs that are known to be affected by lupus and of these only the lungs showed interstitial and peribronchial inflammation (**Fig. 4E** **and Supplementary Fig. 6**). The percentage of TghIL23A mice with pathological findings in their lungs increased in older mice. At 10 weeks of age only 33% of the mice showed lung inflammation while at 20 weeks of age 57% and at 30 weeks of age 73% were affected in their lungs (**Fig. 4E**).

Interestingly, the skin histopathological findings and the local inflammatory milieu together with the pathological manifestations in the lungs bare similarities to the pathological features observed in human cutaneous lupus patients (16,29,31–34).

### 3.5 Human IL23A blockade acts therapeutically and ameliorates TghIL23A cutaneous lupus

The response and dependency of the TghIL23A lupus pathology on the expression of the human IL23A subunit was further studied by treating the mice with guselkumab, a monoclonal antibody that specifically targets and blocks this subunit.

Age and sex matched groups of TghIL23A mice with similar body weights, skin pathology and proteinuria, were treated for 10 weeks, starting from 10 weeks of age, with either 5 or 15mg/kg guselkumab that was administered subcutaneously twice weekly. Clinical monitoring indicated that both doses of guselkumab could significantly reduce the severity of the skin pathology, while only the highest dose could significantly reduce the ear thickness (**Fig. 5A**). Nevertheless, skin and ear histopathological scores were significantly reduced by both doses (**Fig. 5B**).

**Figure 5:**
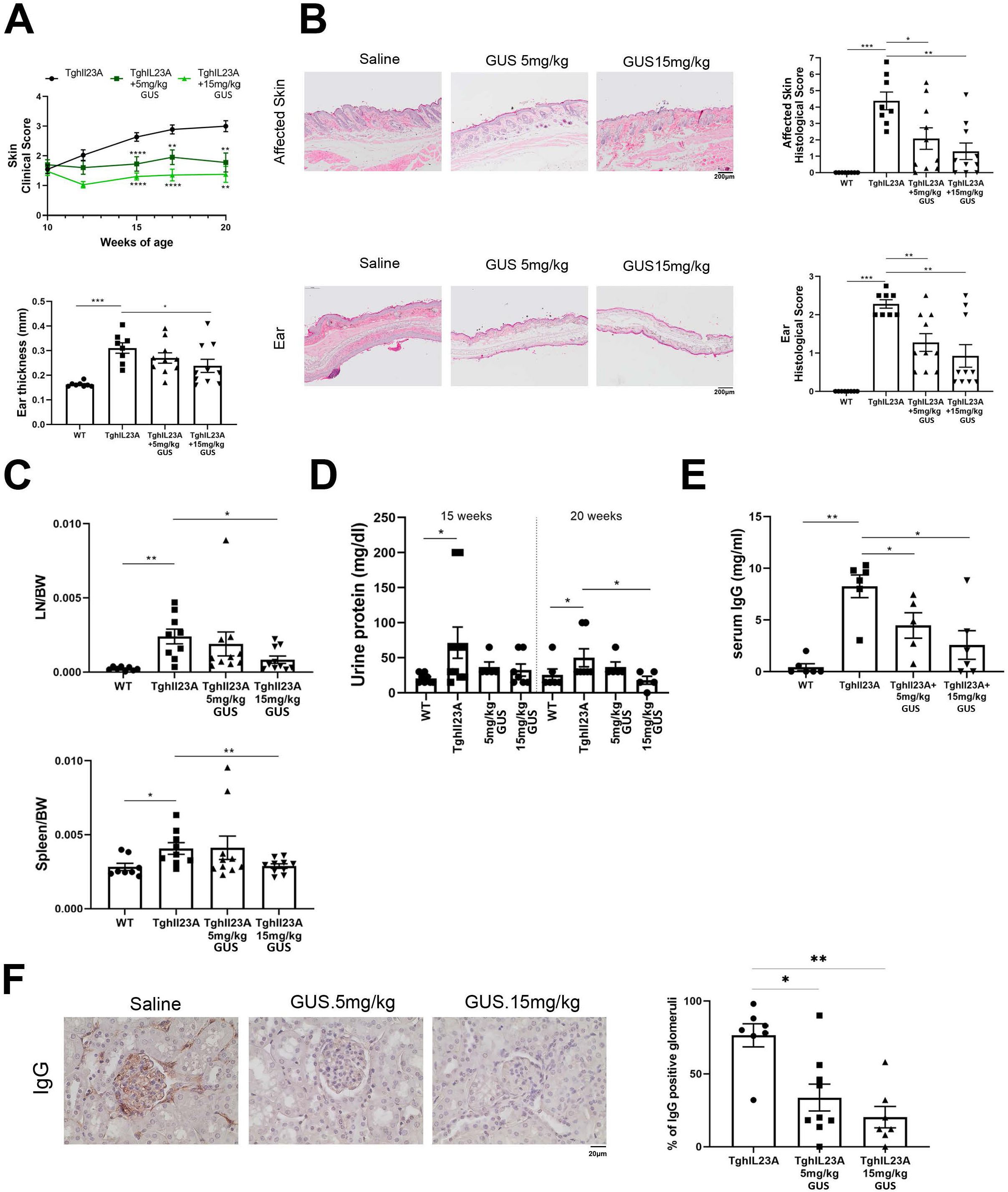
Dose dependent amelioration of the TghIL23A lupus pathology following IL23A blockade. Reduced skin (upper panel) and ear (lower panel) clinical **(A)** and histopathological signs **(B),** as well as reduced LN/BW and Spleen/BW ratios **(C)**, urine protein levels **(D)** IgG circulating levels **(E)** and glomerular IgG deposits **(F)** following a 10-week-long treatment of TghIL23A mice with anti-human IL23A (Guselkumab:GUS). Scale bar 20μm. Data are presented as mean ± SEM; \**P* < 0.04; \*\**P* < 0.0092; \*\*\**P* = 0.0002; \*\*\*\**P* < 0.0001.

Similarly, treatment with the highest dose of guselkumab restored the size of the draining lymph nodes and the spleens of the TghIL23A mice to normal levels (**Fig. 5C**) and returned the urine protein to wild type levels (**Fig. 5D**), while both the 5 and 15mg/kg doses of guselkumab also efficiently reduced the levels of circulating IgGs (**Fig. 5E**).

Finally, treatment with guselkumab efficiently reduced the percentage of IgG positive glomeruli in the kidneys of TghIL23A mice. More specifically, while untreated transgenic mice presented with 76% IgG positive glomeruli, following treatment with 5mg/Kg guselkumab only 34% of the glomeruli stained positive for IgG and following treatment with 15mg/Kg guselkumab this percentage dropped to 20% (**Fig.5F**).

## Discussion

The TghIL23A transgenic mice were generated with the aim of developing a novel mouse model to study the role of deregulated human IL23A expression in pathogenesis. The implication of IL23A deregulation in pathogenic mechanisms has been previously supported by the phenotypes developed in mice lacking the RNA-binding protein tristetrapolin (TTP), a postranscriptional regulator of both TNFα and IL23A. TTP knockout mice spontaneously develop systemic autoimmunity, with features of arthritis, cardiac valvulitis, dermatitis and renal pathology, accompanied by high titers of anti-ds-and ss-DNA autoantibodies (35–37). Since transgenic mice expressing deregulated hTNF were shown to develop arthritis and cardiac valvulitis (38), we questioned whether the deregulated expression of human IL23A could similarly lead to the development of the remaining phenotypes occurring in the TTP-/-mice.

TghIL23A mice were shown to exhibit an overt skin phenotype that included signs of alopecia, as well as the development of erythema and eczematous lesions and with histopathological findings of acanthosis and inflammatory cell infiltration, that are typical signs of dermatitis. Cytokine analysis revealed the local upregulation of IL17A that is consistent with the expected activation of the IL23/IL17 axis in these mice (39,40). Additionally, increased levels of TNF, IL1β and IFNγ cytokines commonly associated with skin inflammation were observed (39,41).

While the skin pathology developing in TghIL23A mice shares several similarities with IL23/IL17-driven psoriasis (39), it also exhibits non-typical features of psoriasis, such as vacuolar interface dermatitis, orthokeratosis, epidermal thinning, epidermolysis and the localized accumulation of IgG deposits, features that resemble those observed in human patients and mouse models of autoimmune skin disorders (24,34). Furthermore, the presence of B220^+^ cells infiltrating the inflamed skin, as well as the enlarged local lymph nodes and spleens, suggest the involvement of an autoimmune component in the TghIL23A driven pathology.

Several lines of evidence support the pathogenic involvement of IL23A in systemic lupus erythematosus (SLE). Specifically, high serum levels of IL23 have been detected in SLE patients and IL23 receptor deficient lupus-prone mice (B6/lpr) are protected from lymphoproliferation, anti-dsDNA antibody production and nephritis (9,42). Additionally, treatment of lupus-prone mice with anti-IL23 antibodies has been shown to ameliorate lupus nephritis (10).

In agreement to the above evidence, our work revealed great similarities between the clinical and histopathological findings of TghIL23A mice to those observed in human SLE patients (11,34). More specifically, the presence of skin pathology, of inflammatory kidney infiltrates, and of interstitial/ peribronchial lung inflammation, as well as the increased circulating levels of anti-nuclear autoantibodies and the local deposition of immunoglobulins in tissues, are key SLE pathology features. Importantly, these pathology findings were shown to be T and B cell-dependent, as they were all ameliorated in a Rag1-/-context, further confirming the involvement of an autoimmune component in the pathology of TghIL23A mice.

IL23A may partner either with the IL23/IL12B subunit to form the IL23 cytokine or with the Ebi3 subunit to form the IL39 cytokine, both of which have been linked to SLE pathogenesis (1,43). Our work revealed that the SLE-related pathology of TghIL23A mice was abolished when crossed with IL23/IL12B deficient mice, therefore confirming that hIL23A exert its pathological effect through dimerization with IL23/IL12B and the formation of the IL23 cytokine.

The TghIl23A pathology, closely resembles that of the Mrl/lpr mouse, one of the most commonly used models of SLE, with shared similarities including the features of cutaneous involvement, splenomegaly, lymphadenopathy, autoantibody formation and renal pathology. Nonetheless, the TghIL23A model is differentiated from the Mrl/lpr model in that it exhibits an extended lifespan due to the slower-developing renal pathology thus offering an extended time frame for the study of the disease and its response to treatment. Additionally, the TghIL23A model allows the direct evaluation of human biologics targeting hIL23A as we have shown that treatment with the anti-hIL23A targeting biologic guselkmab ameliorates the disease. Furthermore, this could be a useful model for testing therapeutic approaches targeting additional components of the IL23 pathogenic pathway.

Overall, with the generation of the TghIL23A mice, we show that the human IL23A subunit dimerizes with the mouse IL23/IL12B subunit, leading to the development of a lupus-like multiorgan autoimmune disease, characterized by autoantibody production and the formation of inflammatory lesions primarily in the skin, kidneys, and lungs. This novel mouse model provides the proof-of-concept for the etiopathogenic role of IL23A in lupus and can serve as a valuable tool for studying the mechanisms underlying autoimmunity and for testing novel therapeutic approaches.

## Acknowledgements

The authors thank Marianna Ragkousi for her help in processing and staining histological samples and Philippos Charalampous for his help in TghIL23ARag1^-/-^ mice breeding and maintenance. We acknowledge support of this work by the InfrafrontierGR/Phenotypos Infrastructure (MIS 5002135), funded by the Operational Programme “Competitiveness, Entrepreneurship and Innovation” (NSRF 2014-2020) co-financed by Greece and the European Union (ERDF), and especially BSRC Fleming’s Transgenics and Flow Cytometry facilities.

## Author contributions

GK, ECV, NK and MCD designed the study and interpreted the experimental results. ECV, CG, LI, and LN performed the experiments and data analysis. ECV, NK and MCD wrote the first draft of the manuscript and all authors were involved in critically revising the final manuscript. All authors reviewed and approved the final manuscript.

