## Supplementary figures and images for "A Novel human IL-23A Overexpressing Mouse Model of Systemic Lupus Erythematosus"

### Supplemental Figures

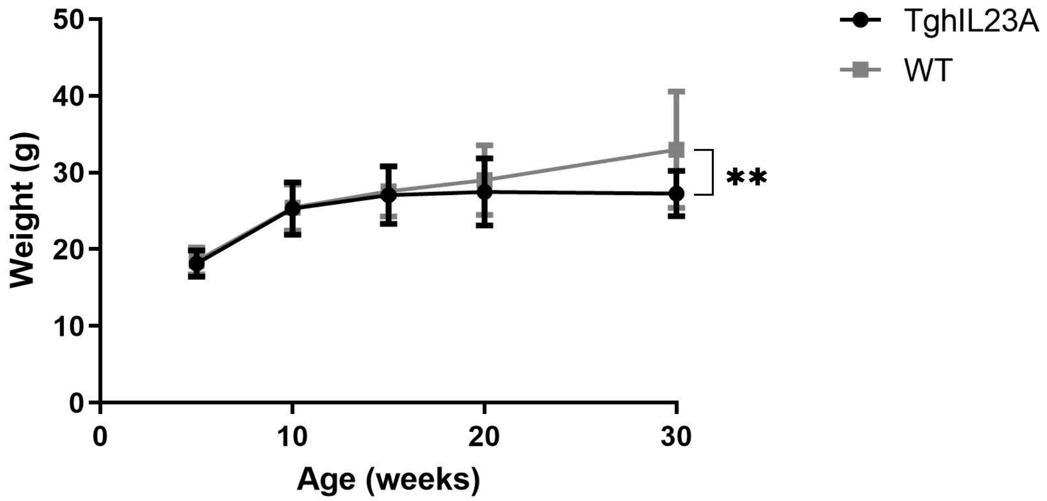


Fig.S1

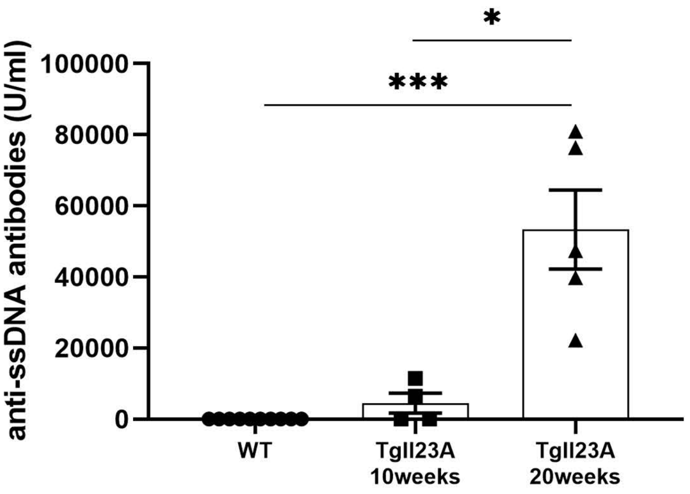

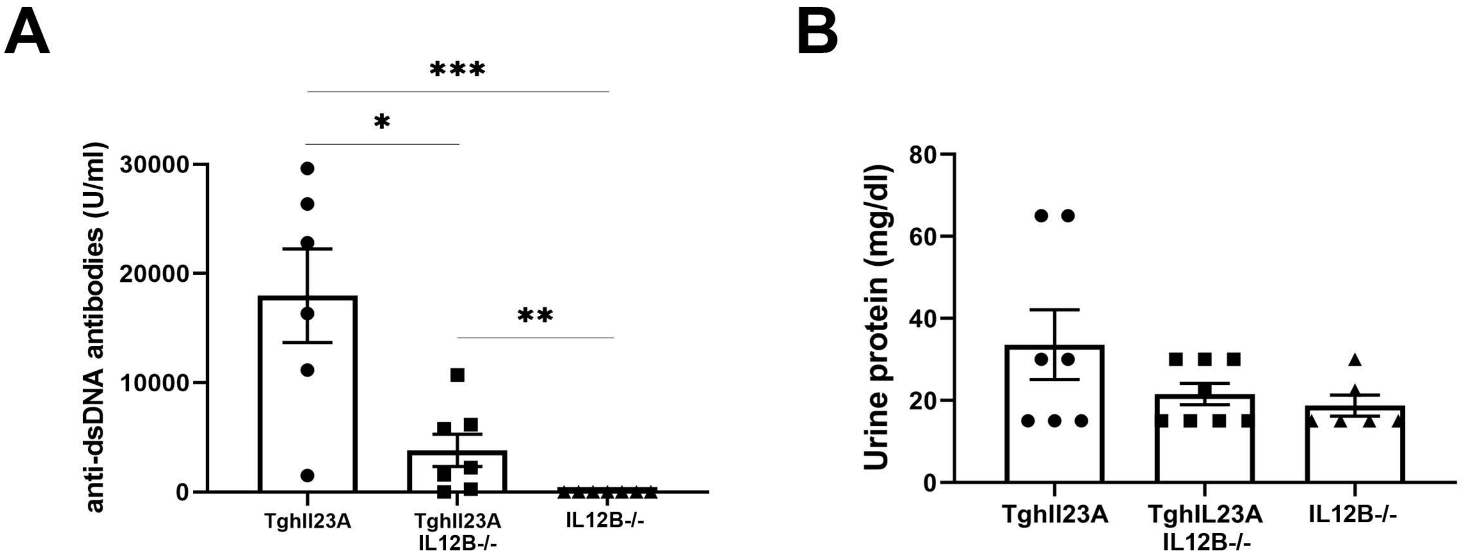

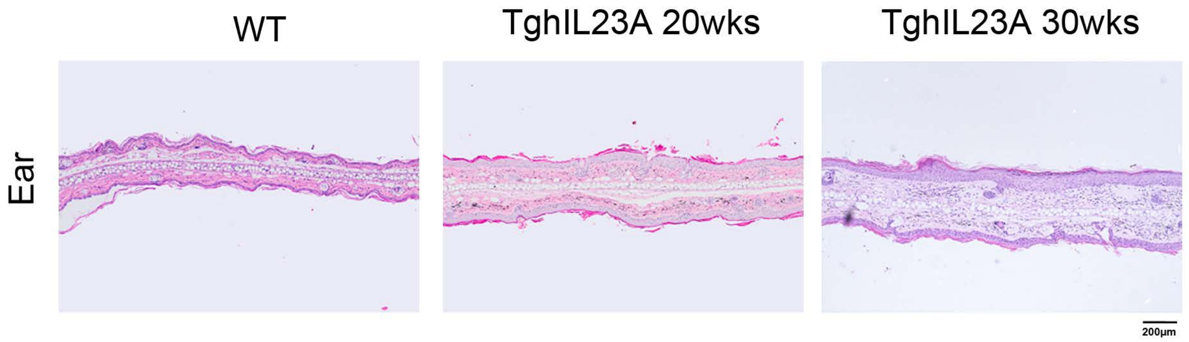

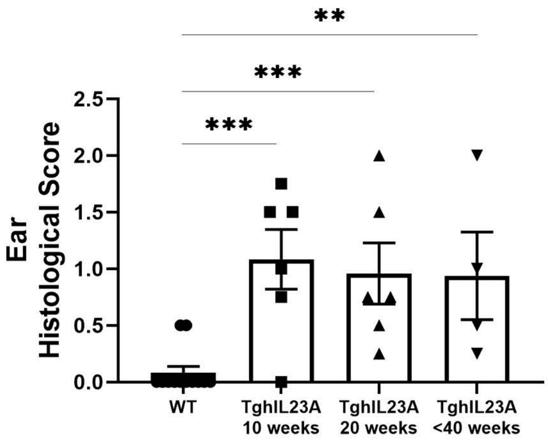

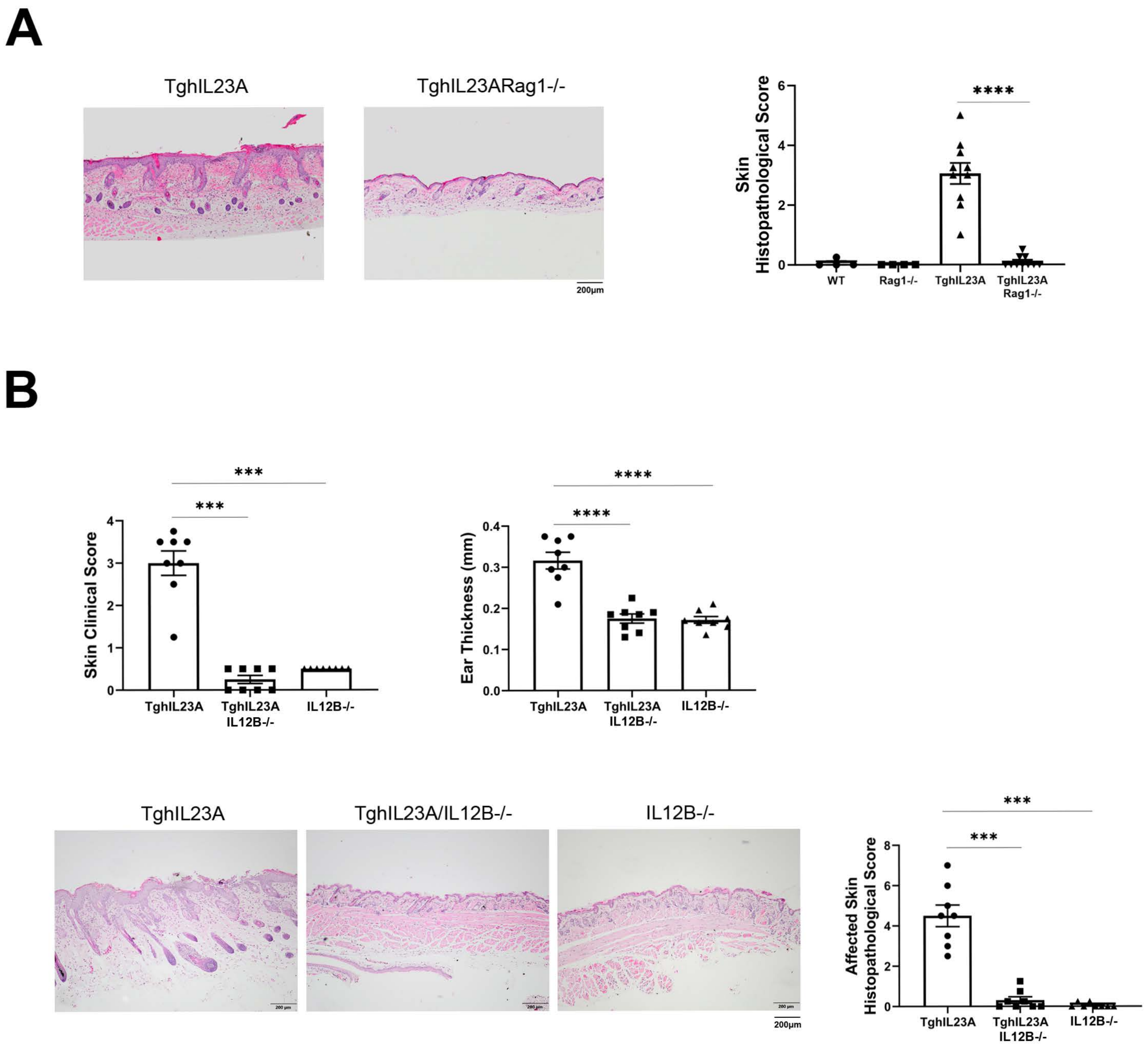

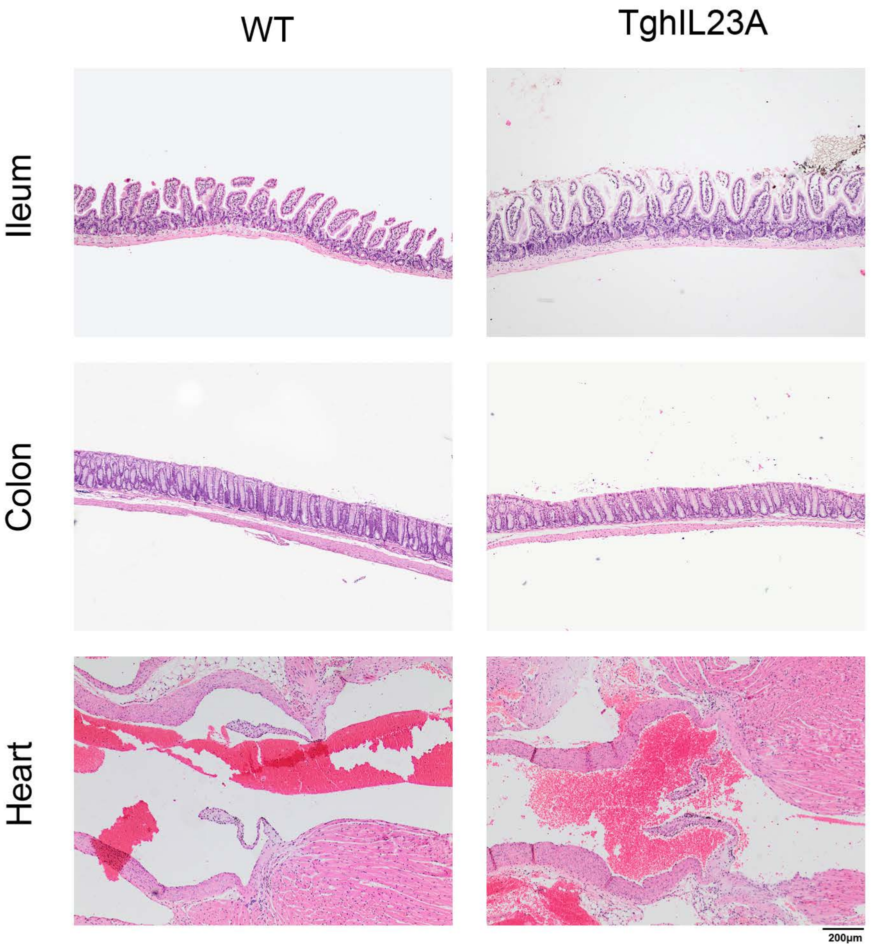


Fig.S6
